# Examination of the contribution of Nav1.7 to axonal propagation in nociceptors

**DOI:** 10.1101/2021.03.12.435114

**Authors:** George Goodwin, Sheridan McMurray, Edward B Stevens, Franziska Denk, Stephen B McMahon

## Abstract

Nav1.7 is a promising drug target for the treatment of pain because individuals with Nav1.7 loss-of-function mutations are insensitive to pain and do not have other serious neurological deficits. However, current peripherally restricted Nav1.7 inhibitors have not performed well in clinical pain trials, which may reflect a lack of understanding of the function of Nav1.7 in the transmission of nociceptive information. Although numerous studies have reported that Nav1.7 has a moderate role in peripheral transduction, the precise contribution of Nav1.7 to axonal propagation in nociceptors is not clearly defined, particularly for afferents innervating deep structures.

In this study, we examined the contribution of Nav1.7 to axonal propagation in nociceptors utilising sodium channel blockers in *in vivo* electrophysiological and calcium imaging recordings from L4 in the mouse. Using the sodium channel blocker TTX (1-10μM) to inhibit Nav1.7 and other TTX-S sodium channels along the sciatic nerve, we first showed that around 2/3^rds^ of nociceptive neurons innervating the skin, but a lower proportion innervating the muscle (45%), are blocked by TTX. In contrast, nearly all large-sized A-fibre cutaneous afferents (95-100%) were blocked by axonal TTX. Characterisation of TTX resistant cutaneous nociceptors revealed that many were polymodal (57%) and capsaicin sensitive (57%).

Next, we examined the role of Nav1.7 in axonal propagation in nociceptive neurons by applying the selective channel blocker PF-05198007 (300nM-1μM) to the sciatic nerve between stimulating and recording sites. 100-300nM PF-05198007 blocked propagation in 63% of C-fibre sensory neurons, whereas similar concentrations did not affect propagation in rapidly conducting A-fibre neurons. We conclude that Nav1.7 has an essential contribution to axonal propagation in only around 2/3^rds^ of nociceptive C-fibre neurons, and a lower proportion (≤45%) of nociceptive neurons innervating muscle.

## Introduction

Non-selective pharmacological blockade of voltage gated sodium channels (Navs) with local anaesthetics, such as lidocaine, at the peripheral or central component of sensory neurons is effective in preventing the transmission of sensory input to the central nervous system (CNS) [15; 23]. In minor surgical procedures, complete pain-relief can be achieved in individuals by injecting local anaesthetics around the appropriate peripheral nerve fibres, and even in extreme pathological conditions, such as neuropathic pain caused by trauma or diabetes, similar results can be achieved [19; 48]. However, as Navs are widely expressed throughout the peripheral and central nervous systems [55], and are essential for electrical signalling in other tissue types, such as cardiac tissue [36], the use of local anaesthetics outside of topical application (e.g. Lidoderm patches in postherpetic neuralgia [16]) and acute nerve block for pain relief is limited.

There are 9 different Nav alpha subtypes (Nav1.1-1.9), five of which are expressed in adult mammalian sensory neurons: the tetrodotoxin sensitive (TTX-S) subtypes Nav1.1, Nav1.6, Nav1.7 and the TTX resistant (TTX-R) subtypes Nav1.9 and Nav1.9 [55]. Nav1.1 and Nav1.6 are predominantly expressed in large sized myelinated A-fibre neurons, which principally encode non-nociceptive information, whereas Nav1.7-Nav1.9 are highly expressed in small unmyelinated C-fibre neurons [55], which are nearly all classed as nociceptors [13]. In theory, selective inhibition of the nociceptive sodium channel subtypes should be effective in providing analgesia in the absence of extensive side-effects because their expression is mostly restricted to DRG neurons [55]. Nav1.7 is the most promising nociceptive subtype to target with an inhibitor because individuals with Nav1.7 loss-of-function mutations are insensitive to somatic and visceral pain [9; 17]. Additionally, affected individuals do not suffer from other sensory abnormalities (except anosmia) [51], and therefore Nav1.7 inhibition might not be expected to produce serious side effects. Considering these findings and observations, it is unsurprising that numerous Nav1.7 blockers are being developed as analgesics. However, many of the small molecule inhibitors developed to date have not been further progressed for the treatment of pain due to lack of efficacy in clinical trials [25].

Most of the Nav1.7 blockers that have failed in the clinic are peripherally restricted (poorly penetrate the blood brain barrier (BBB)) which reduces the likelihood of off-target inhibition of sodium channels in the brain. This means that peripherally restricted Nav1.7 inhibitors are only capable of targeting channels located in the periphery, i.e. the peripheral terminals, axons or DRG, and not the central terminals within the spinal cord, which are physically located across the blood-spinal cord barrier. At the peripheral terminals of cutaneous afferents, pharmacological inhibition of Nav1.7 reduces, but does not abolish, responses to noxious stimuli in around 2/3^rds^ of nociceptors [20; 29], whereas inhibition at visceral afferent terminals is not reported to affect responses to noxious stimulation [14; 20]. The consequences of Nav1.7 inhibition along peripheral axons are less well defined because studies examining the role of the channel in axonal propagation are limited due to the lack of selective blockers. Moreover, the available aryl sulphonamide Nav1.7 blockers are only selective in certain species [1]. A previous pharmacological study in the rat reported that Nav1.7 is essential for propagation in all C-fibre neurons [41]. However, this is in contrast to numerous studies reporting that propagation in a proportion (75-96%) of C-fibres is maintained when Nav1.7 (and other TTX-S channels) are blocked following axonal TTX application [24; 27; 37; 45; 52]. Studies employing genetic knockdown of Nav1.7 also report that a proportion of C-fibres are capable of propagation without the channel [21; 33].

To determine the proportion of nociceptive neurons dependent on Nav1.7 for axonal propagation, we used in vivo electrophysiology and calcium imaging to measure evoked activity before and after application of Nav blockers to peripheral nerve between stimulating and recording sites. Initially, we sought to quantify the proportion of the nociceptive afferents innervating cutaneous and deep structures that are dependent on TTX-S Navs for axonal propagation. Topical application of TTX between stimulating and recording sites revealed a difference in the proportion of nociceptive neurons dependent on TTX-S for propagation between skin and muscle innervating afferents. Further experiments utilising a Nav1.7 selective blocker revealed that Nav1.7 is the major TTX-S channel that is essential for axonal propagation in sensory C-fibre afferents.

## Materials and Methods

### Animals

Adult C57BL/6J mice (n = 30; Charles River, UK) weighing 24-30 g were used for in vivo imaging experiments. Adult male CD1 mice (n = 23; Charles River, UK) weighing 30-45 g were used for electrophysiological experiments. Mice were housed on a 12/12 h light/dark cycle with a maximum of 5 mice per cage, with food and water available *ad libitum*. All experiments were performed in accordance with the United Kingdom Home Office Animals (Scientific Procedures) Act (1986).

### Administration of AAV9-GCaMP6s

We utilised the genetically encoded calcium Indicator GCaMP6s for imaging sensory neuron activity [7]. GCaMP6s was delivered to sensory neurons via a adeno-associated viral (AAV) vector, which was administered to mice pups at P3-P6 as previously described [50]. Briefly, groups of 3-4 mice were separated from their mother and anaesthetised with 0.5-1% Isoflurane. 5 μl of virus AAV9.CAG.GCaMP6s.WPRE.SV40 (Addgene, USA) was injected intraplantar into the left hind paw using a 10μL Hamilton syringe with a 34G needle attached. In experiments examining axonal propagation in muscle afferents the virus was injected subcutaneously in the nape of neck. Following the injection, pups were maintained on a heating pad until they woke up and were then returned to their mother. Mice were separated from their mother after weaning and then were used for in vivo imaging from 7-8 weeks after the injection.

### In vivo imaging of sensory neuron activity using GCaMP

Mice were anesthetized using a combination of 1-1.25 g/kg 12.5% w/v urethane administered intraperitoneally in addition to 0.5-1.5% isoflurane delivered via a nose cone. Body temperature was maintained close to 37°C using a homeothermic heating mat with a rectal probe (FHC). An incision was made in the skin on the back and the muscle overlying the L3, L4, and L5 DRG was removed. Using fine-tipped rongeurs the bone surrounding the L4 DRG was carefully removed in a caudal-rostral direction. Bleeding was prevented using gelfoam (Spongostan^™^; Ferrosan, Denmark) or bone wax (Lukens). The DRG was washed and kept moist using 0.9% saline. The position of the mouse was varied between prone and lateral recumbent to orient the DRG in a more horizontal plane. The exposure was then stabilized at the neighbouring vertebrae using spinal clamps (Precision Systems and Instrumentation) attached to a custom-made imaging stage. The DRG were covered with silicone elastomer (World Precision Instruments, Ltd) to maintain a physiological environment. Prior to imaging, animals received a subcutaneous injection of 0.25-ml sterile normal saline (0.9%) to keep them hydrated. The mouse was then placed under the Eclipse Ni-E FN upright confocal/multiphoton microscope (Nikon) and the microscope stage was diagonally orientated to optimise focus on the DRG. The ambient temperature during imaging was kept at 32°C throughout. All images were acquired using a 10× dry objective. A 488-nm Argon ion laser line was used to excite GCaMP6s and the signal was collected at 500–550 nm. Time lapse recordings were taken with an in-plane resolution of 512 × 512 pixels and a fully open pinhole for confocal image acquisition. All recordings were acquired at 3.65 Hz. In some experiments, the skin was removed from the gastrocnemius muscle. The exposed muscle belly was covered in gauze-soaked KREBS to prevent it from drying out.

### Electrical stimulation of the peripheral terminals

In imaging experiments, the stimulating electrodes used consisted of an anode that was comprised of 6 x 30G needles soldered to a single piece of wire and a cathode that was a single 30G needle. The 6 x 30G needles were inserted transcutaneously into the dorsal surface of the hind paw, while the other needle was inserted into the skin at base of the ankle. A stimulator was used to deliver square wave current pulses of 15mA and 15mS to activate the terminals of both A- and C-fibre neurons. 12 pulses (3Hz, 4s) were delivered at baseline and periodically post-drug application to assess axonal propagation. In electrophysiological experiments, the peripheral terminals were stimulated with a constant current stimulator (Digitimer, UK) connected to two stimulating pins inserted transcutaneously into the dorsal hind paw. Square wave pulses of up to 450us, 450 mA were used to recruit A-fibre neurons and pulses of up to 1ms, 2mA were used for C-fibres. For all experiments involving electrical stimulation, 0.1Hz stimulation was applied throughout the recording to facilitate PF007 binding [1].

### Thermal stimulation of the peripheral terminals

A Peltier device (TSAII, Medoc) with a 16 x 16 mm probe was carefully placed onto the plantar surface of the hind paw ipsilateral to the DRG being imaged and was maintained in position using a V-clamp. The temperature of the block was increased from a baseline temperature of 32°C to 50°C at a rate of 8 °C/s and maintained for 2 seconds before returning to 32°C at the same rate. For cold stimulation, the temperature of the Peltier was decreased to 0°C at a rate of 4 °C/s and maintained for 2 seconds before returning to 32°C.

### Mechanical or chemical stimulation of the peripheral terminals

Mechanically sensitive afferents were identified by 1) pinching using a pair of serrated forceps, 2) brushing using a cotton bud or, 3) flexion and extension of the leg. At the end of some pharmacology experiments capsaicin cream (10%) was gently applied to the plantar surface of the hind paw using a cotton bud.

### Calcium imaging data analysis

The image analysis pipeline Suite2P: (https://github.com/MouseLand/suite2p) [35] was utilised for motion correction, automatic ROI detection and signal extraction. Further analysis was undertaken with a combination of Microsoft Office Excel 2013, Matlab (2018a) and RStudio (Version 4.02). A proportion (0.4) of the neuropil (the local background of each cell) was subtracted from the fluorescent signal for each corresponding cell. To generate normalised data, a baseline period of fluorescence was recorded for each ROI and changes from this baseline fluorescence were calculated as ΔF/F and expressed as a percentage [8]. Cells were grouped into small to medium (≤750μm^2^) and Large (>750μm^2^) sized neurons. A limitation of this technique is the accurate determination of cell sizes. As we are imaging with the pinhole fully open across multiple planes, cells that are located deeper in the tissue can appear larger due to dispersion of the fluorescence. i.e. small cells located deeper in the tissue may appear medium sized. Therefore, we did not split the size groups further into small and medium sized cells. For experiments examining axonal propagation in neurons innervating the hind paw, neurons were tested for responses to heat, cold and mechanical (pinch) stimuli (in that order). Stringent criteria were used for positive responses to thermal and mechanical stimulation of the terminals. A positive response was taken if the average signal during stimulation was 70% plus 4 SD above the relative baseline fluorescence i.e. the average level of fluorescence immediately prior to the stimulation. For electrical stimulation, a positive response was taken if the average signal during stimulation was at least 50% plus 2 SD above the relative baseline. A neuron was classed as not conducting through the treatment site when the average fluorescent signal fell below this threshold. In most instances, traces of non-conducting neurons were double checked by examining raw Ca^2+^ traces. A positive response to capsaicin was taken if the average fluorescence signal during the 5-minute post-capsaicin period was 50% plus 2 SD greater than the 2-minute baseline period. For experiments examining axonal propagation in muscle afferents, only small to medium sized neurons responding at baseline to just muscle pinch (denoted pinch only) and neurons responding to just hind paw brush (denoted brush only) that were blocked by lidocaine at the end of the experiment were analysed. To facilitate drug block, neurons were activated periodically by brush and pinch stimulation throughout the experiment (See Fig 2A). At the end of each drug application period, the stimulation procedure was performed twice to reduce likelihood of missing receptive fields activated at baseline.

**Figure 1.**
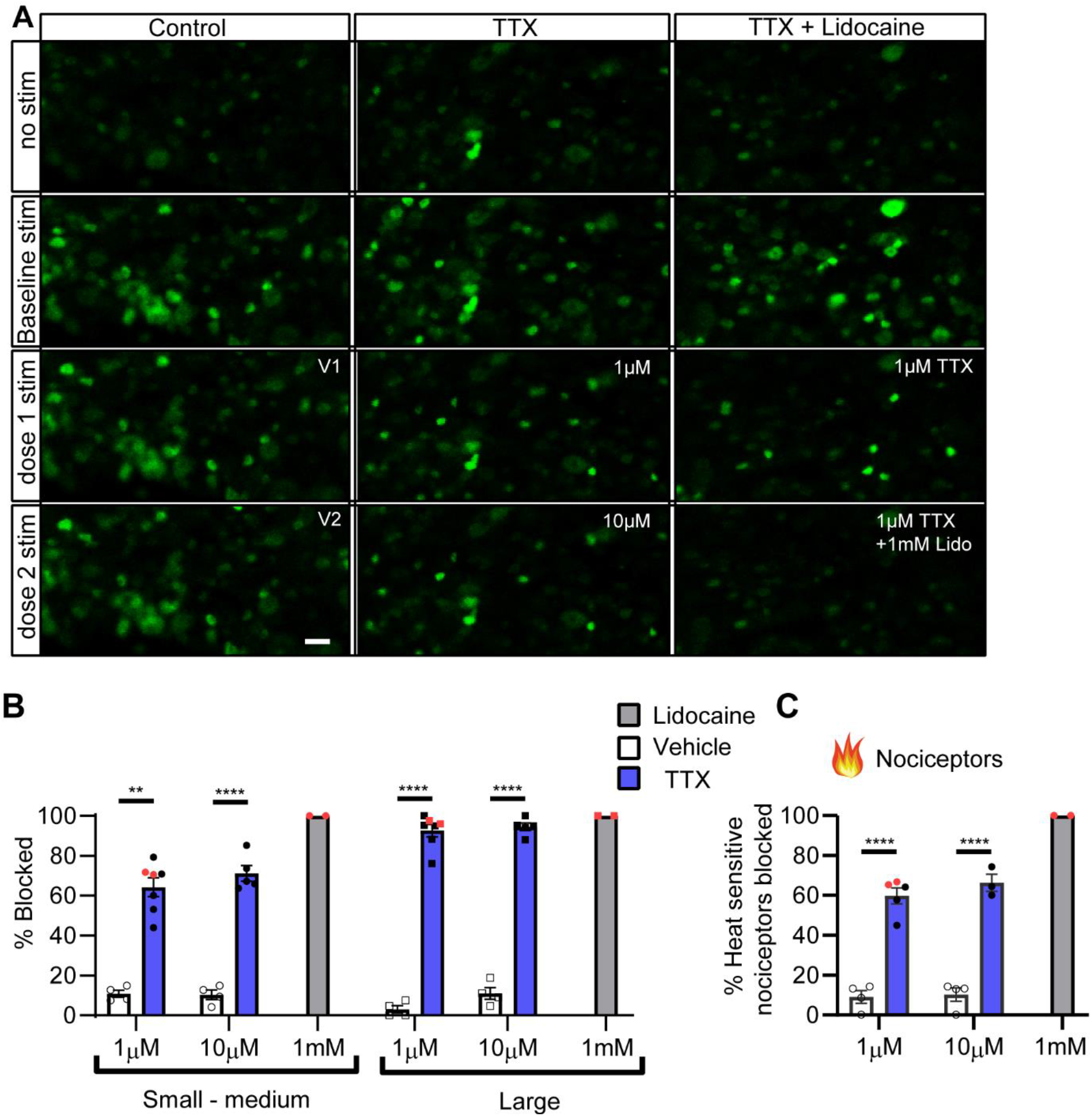
Axonal TTX application blocks propagation in nearly all large sized neurons but only ~2/3 of small to medium sized neurons. Example images of L4 DRG cell bodies from in-vivo GCaMP6s time lapse recordings (A). Frames were extracted at baseline and at the peak of electrical stimulation at baseline and following each drug dose. The bar graphs show the proportion of neurons blocked following drug application to the nerve when neurons are grouped by size (B) and for neurons responding to noxious heat stimulation of the terminal (C). Each point represents the proportion of neurons blocked in an individual animal. 47-136 small to medium, 11-41 large neurons and 14-54 heat responsive neurons were sampled between animals. Note that all sizes of neurons are activated by electrical stimulation of the paw at baseline. The electrically induced response was blocked in nearly all large sized cells following TTX application to the nerve, but only a proportion of small to medium sized neurons (middle panel). Small to medium sized neurons not blocked by TTX were blocked by lidocaine (right panel in A, grey bars in B & C). The red coloured dots denote the experiments in which lidocaine and TTX were applied to the nerve following 1μM TTX. Bar graphs represent mean ± SEM. Scale bar = 50μM. 10x magnification. 3-way ANOVA for B) and RM 2-way ANOVA for C). ** p<0.01, **** p<0.0001 vs control Sidak’s post hoc.

**Figure 2.**
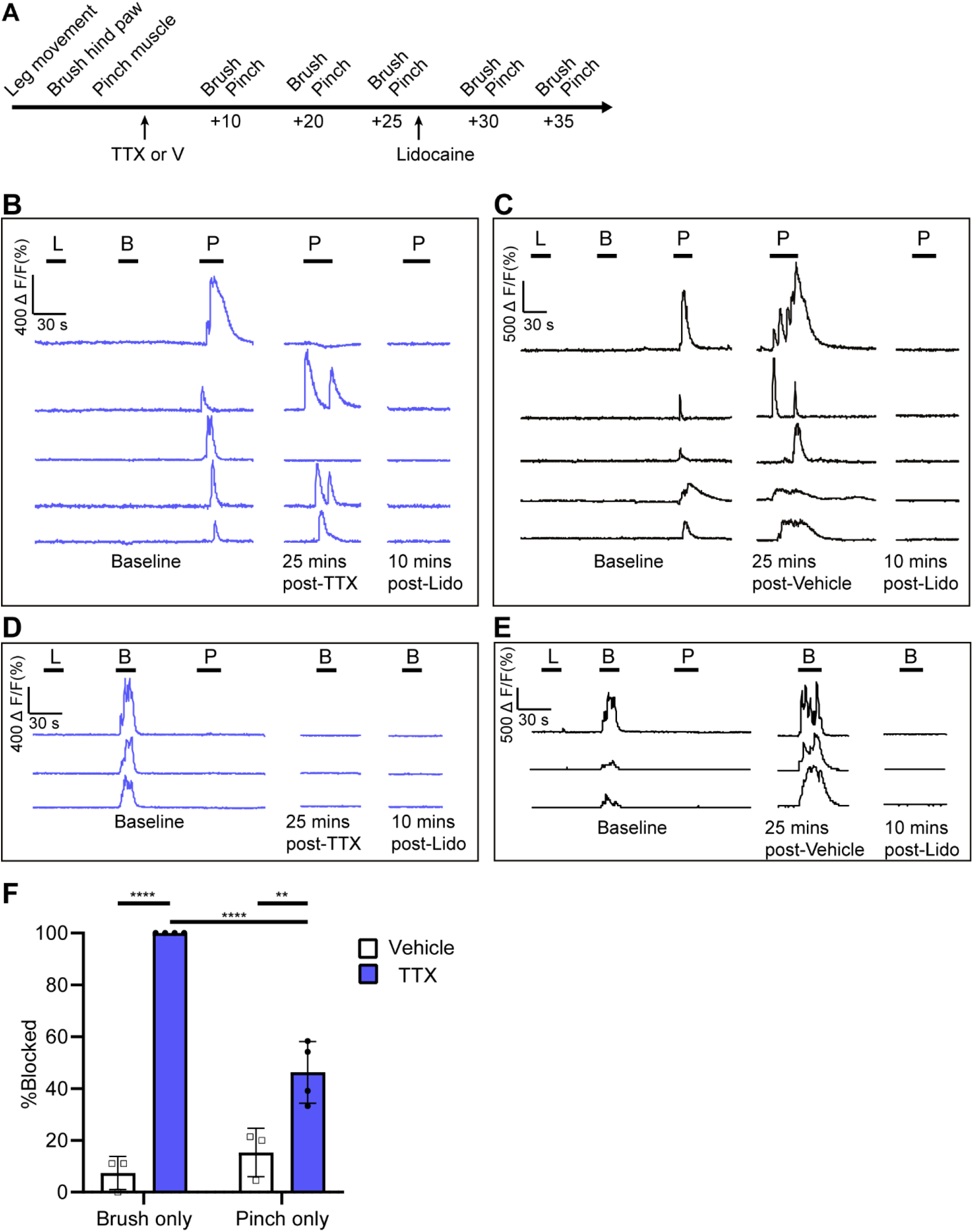
Axonal TTX application blocks propagation in all brush responsive hind paw neurons but only 46% of pinch responsive small to medium sized muscle afferents. Experimental schematic is shown in (A). Example calcium traces are presented in (B-D). Traces are shown for pinch responsive small to medium neurons (B,C) or brush responsive hind paw neurons (D,E) before and after TTX (B,D - blue) and Vehicle (C,E - black) application to the nerve. Note that only neurons responding at baseline to just muscle pinch or just brush that were blocked by lidocaine were analysed. The bar graph shows the average proportion of neurons blocked after 25 mins of drug application for each group (F). Between 6-28 brush and 12-30 pinch neurons were sampled between animals. Horizontal bars represent stimulation periods. L – leg movement, B brush of plantar surface of hind paw, P – pinch of exposed muscle belly. Bar graphs represent mean. 2-way ANOVA ** p<0.01, **** p<0.0001 vs control Sidak’s post hoc. See also Movie 1.

### In vivo electrophysiology

*In vivo* single-unit electrophysiological recordings from A- or C-fibre axons in the L4 dorsal root were carried out in CD1 mice (n = 23), as previously described in the rat [3; 18]. Animals were anesthetised using a combination of 1-1.25 g/kg 12.5% w/v urethane intraperitoneally in addition to 0.5-1.5% isoflurane delivered via a nose cone. The body temperature was maintained at 37°C using a rectal thermistor probe connected to a heated pad (Harvard Apparatus, Kent, United Kingdom). A lumbar laminectomy was performed from L2 to L5 to expose the spinal canal. The surrounding skin was sutured to a metal ring to form a pool filled with warmed mineral oil. The dura mater was opened and the left L4 dorsal root was cut close to the spinal cord. The cut end of the dorsal root was placed onto a glass platform (4 × 2.5 mm).

Fine filaments were teased from the cut end of the dorsal root using finely sharpened forceps and individually placed onto paired gold recording electrodes. Extracellular potentials were band-passed filtered (10-5,000 Hz) and amplified (1k) and digitised using Spike 2 software (Cambridge Electronic Designs, Cambridge, United Kingdom). Mechanical receptive fields were searched for and once a field had been identified, needle electrodes were inserted into the skin at the receptive field location. Electrical stimulation was applied as described above, and the number of fibres on each filament was determined by slowly increasing the stimulus amplitude until the maximum number of waveforms was elicited. To prevent a single electrical stimulus causing >1 action potential to be produced from the same neuron, a cut-off amplitude of 450uA was used for A-fibres and 2mA for C-fibres. Only clearly identifiable waveforms were studied (<8 fibre waveforms per filament). Neurons were classified based on waveform shape and conduction velocity, which was determined from the conduction distance and latency of individual waveforms. Conduction velocities above 16m/s were considered Aα/β-fibres, and below 1.2 m/s were considered C-fibres [56]. Neurons conducting in the Aδ-fibre were identified in some experiments, but we preferentially searched for C-fibres because our primary aim was to compare the effects of Nav1.7 in this population. Once a small (2-7) group of A- or C-fibre neurons with conduction velocities that were stable had been identified, the terminal was stimulated at 0.1Hz and drugs were applied topically to the sciatic nerve. Neurons were considered to not be conducting through the treatment site when they failed to produce an action potential following 6 consecutive electrical stimulations, or, if at the end of the experiment, no longer responded to mechanical stimulation of the receptive field.

### *In vivo* topical drug application

The sciatic nerve was exposed at the level of the midthigh and was freed from surrounding connective tissue. A 2-3 mm portion of the nerve, located 1-2mm proximal to the trifurcation, was carefully desheathed using the tip of a 30G insulin needle. To isolate the desheathed portion from the surrounding tissue, the nerve was covered in 4% agar (MP Biomedicals) dissolved in 0.9% saline. Once it had set, a well (~vol 25-30uL) was cut out around the desheathed portion and was filled with warmed (37°C) Krebs solution. All drugs were warmed (37°C) before application, and the well was periodically replaced with fresh drug solution throughout recordings. Each drug concentration was applied sequentially for a duration of 20-30 minutes.

### Drugs and Chemicals

All drugs were dissolved to make stock solutions and then frozen at −20°C until their day of use. TTX (Acros Organics, Fisher Scientific) and 4,9-Anhydrotetrodotoxin (Tocris, UK) were dissolved to 1mM in 0.2 mM citric acid (pH 4.8). PF-05198007 (a gift from Dr S McMurray [1]), was dissolved in DMSO to 10mM. Lidocaine hydrochloride (Sigma, UK) was dissolved in 0.9% w/v sodium chloride to a concentration of 74mM (2% w/v). Drugs were diluted down to their final concentrations in either freshly carbogenated (95% O2 5% Co2) Krebs solution (In mM: NaCl, 118; KCl, 4.7; NaHCO3, 25; KH2PO, 1.2; MgSO4, 1.2; glucose, 11; and CaCl2, 2.5) adjusted to pH 7.4.

### Quantification and Statistical analysis

Graphing and statistical analysis was undertaken with a combination of Microsoft Office Excel 2013 and GraphPad Prism (Version 8). All data sets were checked for normality using Kolmogorov-Smirnov tests. Details of statistical tests and sample sizes are recorded in the appropriate figure legends. Main effects from ANOVAs are expressed as an F-statistic and P-value within brackets. All data plotted represent mean ± SEM. Statistical differences between groups in *in vivo* imaging experiments involving different drug groups, neuron sizes and doses were compared using a 3-way repeated measures analysis of variance (ANOVA) with Sidak’s post hoc comparisons. For imaging experiments comparing drug effects on propagation in heat responsive nociceptors, statistical differences were determined using a 2-way repeated measures ANOVA with Sidak’s post hoc comparisons. For imaging experiments involving muscle afferents, statistical differences between groups were determined using a 2-way ANOVA with Tukey’s post hoc comparison. The proportion of neurons blocked by TTX in different stimuli response groups was compared using Chi-squared test. For analysis of the *in vivo* electrophysiology data, conduction velocities were compared between models using Kruskal-Wallis tests and the proportion of neurons blocked between drug groups was compared using Fishers exact test.

## Results

### Axonal TTX application has differential effects on propagation depending on fibre type and innervation territory

Electrical stimulation of the plantar surface of the hind paw activated neurons of all sizes (small/medium neurons mean 71 (SEM 8.7) and large mean 22 (SEM 2.3); n = 11 animals; Fig 1A – second panel). The total number of neurons recruited via electrical stimulation varied between experiments (range of 58-157) and therefore the proportion of neurons responding to electrical stimulation at each time point was averaged between animals. In a significant number of neurons, TTX applied to the sciatic nerve between stimulating and recording sites blocked propagation; main effect of TTX: F (1, 7) = 339.6, p<0.0001; 3-Way ANOVA. Main effects of dose (F (1, 7) = 8.58, p<0.05) and neuron size (F (1, 7) = 14.40, p<0.01) were also statistically significant, as were several interactions, including between neuron size x TTX (F (1, 7) = 24.84, p<0.01) – a result in line with expectations, as TTX should preferentially block large sized neurons. Thus, 1μM TTX blocked 61.4% (SEM 6.3) small to medium and 90.9% (SEM 4.2) large sized neurons, which was significantly different to that of the control for each corresponding size group (mean block = 10.9%, SEM 1.8 and 3.0%, SEM 2.9; p<0.01 & p<0.0001 Sidak’s post-hoc comparisons test for small to medium and large sized neurons respectively; n= 4-5 mice group; Fig 1A-left & middle panels & 1B). Subsequent application of 10 μM TTX produced a small and non-significant increase in the number of axons blocked (average of 71.0% (SEM 3.9) and 94.5% (SEM 1.9); p= 0.41 and p=0.85 Sidak’s post-hoc comparisons compared to 1 μM TTX for small to medium and large sized neurons respectively; Fig 1A - middle panel & 1B).

Not all small to medium sized DRG neurons are nociceptors and therefore we examined the effects of TTX on propagation only in the neurons that responded to noxious heat stimulation at the beginning of the experiment. TTX blocked propagation in a significant number of heat sensitive nociceptors; main effect of TTX: F (1, 5) [Drug] = 84.69, p<0.001; Repeated measures 2-Way ANOVA. There was also a small but significant interaction between Dose x Drug group (F (1, 5) = 15.93, p<0.05). The average number of heat sensitive nociceptors blocked by TTX was 55.6% (SEM 5.6) at a concentration of 1 μM and 66.3% (SEM 4.2) at 10 μM, which were both significantly different to their corresponding time-matched controls (mean block = 9.1%, SEM 3.2 and 10.2%, SEM 2.4 following vehicle 1 & 2 respectively; both p<0.0001 Sidak’s post-hoc comparisons; n= 3-4 mice; Fig 1C). The proportion of neurons not blocked by TTX were blocked by subsequent application of 1mM lidocaine/ 1μM TTX to the sciatic nerve (100% block for all groups; n= 102-131 small to medium and 25-51 large neurons averaged from n = 2 mice; Fig 1A – experiments denoted by red dots in right panel, 1B & C).

The proportion of nociceptive neurons innervating muscle afferents sensitive to axonal TTX application was examined (See Fig 2A for experimental design). Presumptive muscle afferent nociceptors were identified as small to medium sized neurons responding at baseline only to strong muscle pinch, but not leg movement or hind paw brush (eliminating low threshold proprioceptive and tactile afferents). TTX block in the presumptive muscle nociceptors was compared to non-nociceptive large sized neurons that only responded to hind paw brush. The total number of neurons activated varied between experiments (6-28 neurons activated by brush only, 12-30 neurons activated by pinch only) and therefore the proportion of baseline responding neurons also responding at each timepoint was averaged between animals. 1 μM TTX caused a significant block of propagation in both brush and pinch responsive groups (F [Drug] (1, 10) = 191.2 p<0.0001; 2-way ANOVA). There were also significant main effects between brush and pinch groups (F (1, 10) = 26.34 p<0.001; 2-way ANOVA) and the interaction between drug and stimuli groups (F (1, 10) = 47.7, p<0.0001). 1μM TTX blocked 100% of brush afferents and 46.2% (SEM 6.0) pinch afferents, which was significantly different to that of the control for each corresponding group (mean block = 7.4%, SEM 3.7 and 15.3%, SEM 9.4; n=3-4 mice/group; p<0.0001 & p<0.01 Tukey post hoc comparison for brush and pinch afferents respectively; Fig 2 B-F, Movie 1).

### Tetrodotoxin resistant neurons innervating the hind paw are comprised of capsaicin sensitive polymodal nociceptors

Responses to heat, cold and mechanical stimuli were assessed prior to electrical stimulation in most topical drug experiments and neurons were grouped according to their responses to different stimuli (Fig 3A for summary of TTX experiments). The level of polymodality within a neuronal population was assessed by determining the number of thermally responsive neurons also responding to mechanical stimulation and the number of mechanically sensitive neurons also responding to thermal stimulation (Supplementary table 1). Most temperature sensitive neurons also responded to mechanical stimulation (67.1%, SEM 3.4 and 73.9%, SEM 3.5 for heat and cold groups respectively, average from n = 13 mice; Supplementary Fig 1A). In contrast, a lower proportion of mechanically sensitive neurons also responded to heat stimulation (mean = 32.4%, SEM 3.3) and very few also responded to cold stimulation (mean = 5.2%, SEM 0.5; Supplementary Fig 1B). Levels of polymodality were increased when assessed in neurons that responded to electrical stimulation (Supplementary table 1).

**Figure 3.**
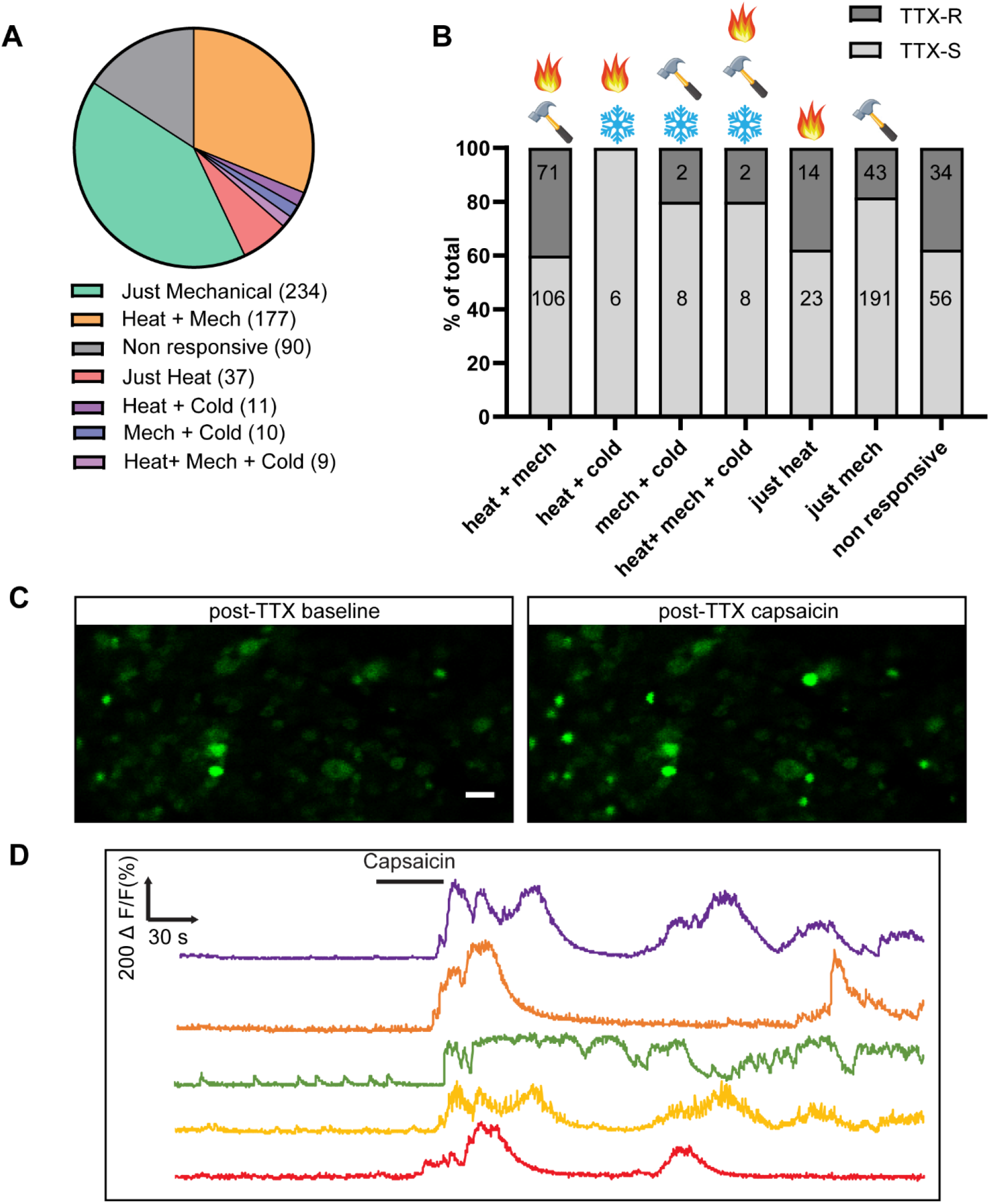
Small to medium sized neurons resistant to axonal TTX are comprised of polymodal and unimodal nociceptors and many are capsaicin sensitive. Pie chart represents electrical responding small to medium sized neurons grouped according to their responses to different stimuli (A). Within each of these neuron groups, the neurons blocked by TTX (TTX-S), or not blocked by TTX (TTX-R) are expressed as a percentage of the total number of neurons within each group (B). 66-136 small to medium sized neurons pooled from n = 6 mice. A proportion of neurons responded to capsaicin following TTX application (C&D). Example images of L4 DRG cell bodies from in-vivo GCaMP6s time lapse recordings (C). Frames were extracted after 1μM TTX application before and 90s after application of capsaicin. Example calcium traces are shown in (D). Example Images were taken at 10x magnification, scalebar = 50 μm. See also Movie 2.

The small to medium sized population not blocked by 1μM axonal TTX application contained a higher proportion of polymodal neurons (56.8%, 75/132 responsive neurons) compared to the population blocked by TTX (37.4%, 128/342 responsive neurons; p<0.001, Chi-square test with Yates’ correction). The polymodal mechano-heat (M-H) group contained the highest proportion (40.1%, 71/177) of neurons that were not blocked by axonal TTX application, whereas the proportion of mechano-cold (M-C) polymodal neurons was lower (20%, 2/10), but not significantly different (p = 0.35 vs M-H group, Chi-square test with Yates’ correction; Fig 3B). All the polymodal heat-cold neurons (6/6) were blocked by axonal TTX application. The proportion of unimodal neurons responding to heat not blocked by axonal TTX was 37.8% (14/37), which was significantly different compared to neurons responding to just mechanical (18.4%, 43/234; p < 0.05, Chi-square test with Yates’ correction; n = 6 mice). As nearly all large sized neurons were blocked by axonal TTX, no comparisons were made within this population. To determine how many TTX-R neurons were TRPV1 positive, 10% capsaicin cream was applied to the corresponding area of the hind paw at the end of the experiment. Application of capsaicin to the hind paw after 1-10 μM TTX application to the nerve caused a positive response in 56.9% (SEM 3.7; n = 3 mice; Fig 4C & D, Movie 2) of neurons that were still conducting, which was greater than the number of responders following control (mean response = 37.5%, SEM 3.0; n = 2 mice).

**Figure 4.**
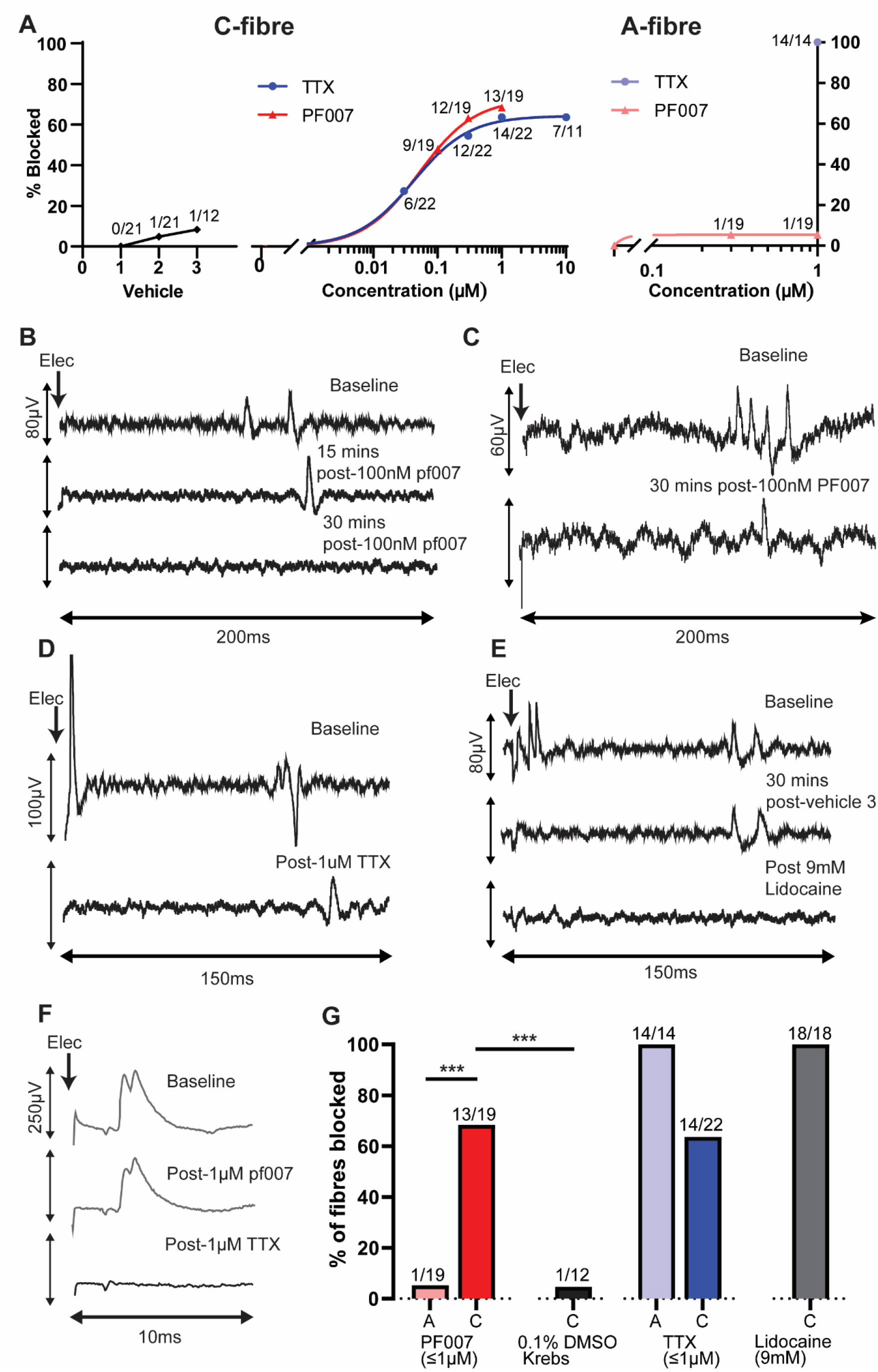
Topical application of PF007 blocked conduction in 2/3rds of C-fibre sensory neurons. 2/3^rds^ of C-fibre sensory neurons were blocked following topical application of PF007 or TTX (A – middle panel, B, C & D), but not following vehicle (A-left panel, E). A-fibre neurons were not blocked by PF007 but were blocked subsequently by application of TTX (A – right panel, F). Example traces of C-fibre (B-E) and A-fibre (F) neurons on electrical stimulation of the hind paw. Conduction slowing typically preceded block following application of PF007 (B & C) and TTX (D), whereas slowing was not apparent following vehicle application (E) or following PF007 application to A-fibre neurons (F). (G) Summary of electrophysiology data *** p<0.001 Fishers exact test. n=5-6 mice/group. Number of neurons in each group shown in (A). Note that the final application of vehicle was not possible in n=2 mice due to technical issues.

### Nav1.7 is the major TTX-S subtype in the proportion of nociceptors blocked by TTX

*In vivo* recordings were made from 19 A-fibre (n=5 mice) and 62 C-fibre sensory neurons (n=18 mice) that had conduction velocities which ranged from 19.9-40.3 m/s and 0.4-1.1 m/s respectively. Conduction velocities of C-fibre neurons at baseline were not significantly different between the three treatment groups (p= 0.26, Kruskal-Wallis test; Table 1). Mechanical receptive fields were identified in the majority of A-fibre and C-fibre neurons identified electrically and were all located on the plantar surface of the hind paw (Table 1). C-fibres with mechanical receptive fields only responded to strong mechanical stimulation of the hind paw indicating that they were nociceptive.

**Table 1.** Properties of A- and C-fibre neurons identified in experiments.

|  | A-fibre | C-fibre |  |  |
| --- | --- | --- | --- | --- |
|  | PF007/TTX | Vehicle/Lidocaine | PF007 | TTX |
| CV (m/s), median (IQR) | 21.72 (6.27) | 0.54 (0.09) | 0.52 (0.10) | 0.55 (0.08) |
| No. of neurons (mice) | 19 (5) | 21 (6) | 19 (6) | 22 (6) |
| No. of mechanical receptive fields identified | 13 | 12 | 11 | 12 |

We utilised the selective Nav1.7 blocker PF007 to examine how many nociceptors depend on Nav1.7 for propagation[1]. Preliminary dose range finding studies carried out in *ex vivo* compound action potential (CAP) experiments found that 100-300nM PF007 applied between stimulating and recording sites reduced the C-CAP but not the A-CAP (data not shown). In *in vivo* recordings, topical application of 100nM-1μM PF-007 blocked conduction in 68.4% (13/19) of C-fibre neurons, whereas vehicle application only blocked conduction in 8.3% (1/12; p<0.001 Fishers exact test; Fig 4 A & G). Notably, the majority of C-fibre neurons blocked by PF007 were blocked at the most selective concentration of PF007 (47.4% blocked following 100nM PF007; Fig 4B & C). Application of TTX blocked conduction in a similar proportion (63.6%) of C-fibre neurons (14/22 blocked following 30nM-1μM; Fig 4A & D). Further application of 10μM TTX did not cause any additional increase the proportion of fibres blocked (7/11 blocked following 30nM-10μM from 2 mice). Conduction slowing typically preceded conduction block in C-fibre neurons following PF007 and TTX application, but not vehicle (See examples in traces 4B, D & E). Application of 9mM Lidocaine blocked conduction in all C-fibre neurons (18/18 from n=5 mice; Fig 4E & G). In comparison to C-fibres, the application of 300nM-1μM PF007 did not cause significant conduction block in A-fibre neurons (5.3% (1/19, p<0.001 Fishers exact test compared to C-fibre; Fig 4A, F & G). All A-fibre neurons that were not blocked by 300nM-1uM PF007 were blocked subsequently by application of 1μM TTX (14/14 from n = 4 mice, Fig 4F & G).

Topical drug experiments with PF007 were repeated in GCaMP6s labelled mice to corroborate electrophysiological findings. Surprisingly, in this preparation, application of PF007 to the sciatic nerve between stimulating and recording sites did not exert a significant block on axonal propagation (F (1, 6) [Drug] = 0.58, P=0.47; 3-Way ANOVA), even when the concentration was increased from 300 nM to 1 μM (F (1, 6) [Dose] = 2.55, P=0.16; 3-Way ANOVA). There was, however, a significant main effect of neuron size (F [Size] (1, 6) = 19.44, p<0.01); the proportion of small to medium sized neurons blocked at 300 nM (17.3%, SEM 4.1) and 1 μM (21.1%, SEM 5.0) was significantly greater than that of the large sized neurons at each corresponding dose (300 nM mean block = 3.8%, SEM 2.5; 1 μM mean block = 2.6%, SEM 2.6; p<0.05 and p<0.01 respectively, Sidak’s post-hoc comparison; n = 4 mice/group; Fig 5B). Similar to the % block in the small to medium sized group, PF007 only blocked 17.9% (SEM 5.6) and 15.0% (SEM 5.5) of heat sensitive nociceptors at 300 nM and 1 μM respectively, which was not significantly different to that of the control (mean block = 9.1%, SEM 3.2 and 10.2%, SEM 3.4 following vehicle 1 & 2 respectively; F (1, 6) [Drug] = 1.16 P=0.32, 2-way ANOVA; Fig 5c).

**Figure 5.**
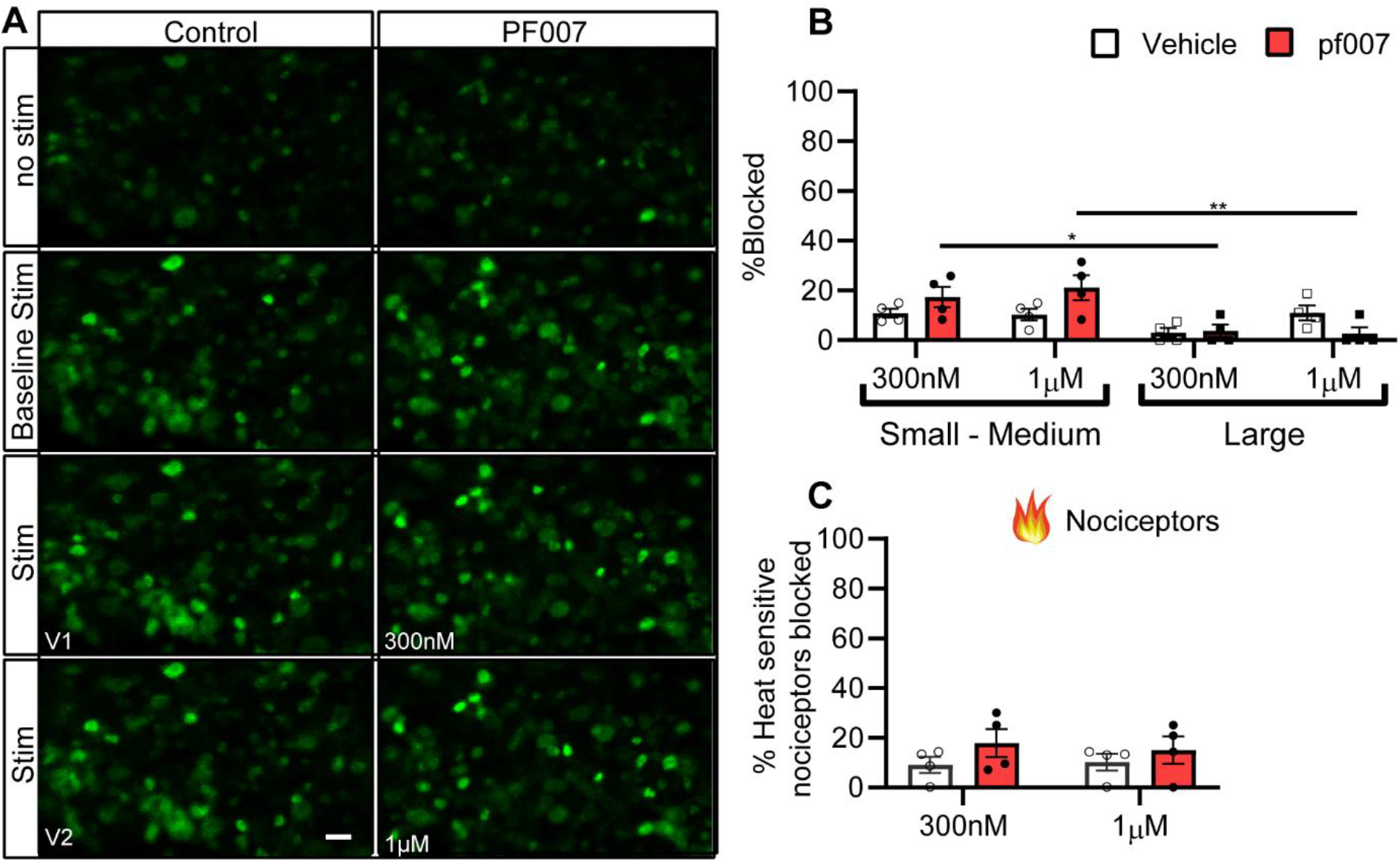
Axonal PF007 application had minimal effects on propagation when assessed in mice via GCaMP6s in vivo imaging. Example images of L4 DRG cell bodies from in-vivo GCaMP6s time lapse recordings (A). Frames were extracted at baseline and at the peak of electrical stimulation at baseline and following each drug dose. The bar graphs show the proportion of neurons blocked following drug application between stimulating and recording site when neurons are grouped by size (B) and for neurons responding to noxious heat stimulation of the terminal (C). Each point represents the proportion of neurons blocked in an individual animal. Between 31-124 small to medium, 6-29 large neurons and 14-27 heat responsive neurons were sampled between animals. The electrically induced response was blocked in few neurons after topical application of PF007 to the sciatic nerve (A-right panel, B & C). Note that these data were collected at the same time as the TTX data in Fig.1 – the vehicle control results are therefore identical. Bar graphs represent mean ± SEM. Scale bar = 50μM. 10x magnification. 3-way ANOVA for B) and RM 2-way ANOVA for C). ** p<0.01 vs control Sidak’s post hoc test.

The reason for the discrepancy in the proportion of nociceptors blocked by PF007 between electrophysiological and calcium imaging experiments was not clear and therefore we decided to use a different class of sodium channel blocker to further investigate this. Due to the limited availability of Nav1.7 selective inhibitors in the mouse, we used 4,9 AH TTX, a selective blocker of Nav1.1 and Nav1.6 [10; 39], and then blocked Nav1.7 in addition to these channels with TTX. The population dependent on Nav1.7 for propagation can be estimated by subtracting the number of neurons blocked after 4,9 AH TTX from the total blocked after subsequent TTX application. In pilot dose range finding experiments, application of 1 μM 4,9 AH TTX did not block propagation in small to medium or large sized neurons (data not shown). In contrast, application of 3 μM 4,9 AH TTX to the sciatic nerve in-between stimulating and recording sites caused propagation failure in a significant number of neurons (F (1, 6) [Drug] = 77.12 P=0.0001). The main effects of Size (F (1, 6) = 104.1, p<0.0001) and the interaction between Drug x Size (F (1, 6) = 31.73, p<0.01) were also significant. 4,9 AH TTX blocked conduction in 30.4% (SEM 5.7) of small to medium sized neurons, which was greater, but not significantly different, to that of the time-matched control (mean block = 16.3%, SEM 1.6; p= 0.23, Sidak’s post-hoc comparison; n = 4 mice/group; Fig 6A & B). In contrast, 4,9 AH TTX blocked (80.8%, SEM 2.6) large sized neurons, which was significantly different to the control (mean block = 8.1%, SEM 2.7%, p<0.0001, Sidak’s post-hoc comparison; Fig 6A & B, Movie 3). Subsequent addition of 1 μM TTX to the sciatic nerve further impacted axonal propagation. The main effect of TTX addition was significant (F (1, 6) = 1136, p<0.0001), as were several interactions, including that between Drug x Size x TTX addition (F (1, 6) = 28.24, p<0.01). The proportion of small to medium neurons blocked by TTX increased to 73.5% (SEM 2.1) in the 4,9 AH TTX group and 70.3% (SEM 4.7) in the vehicle group, which were both significantly different to the block produced previously (both p<0.001, Sidak’s post-hoc comparison to block following 4,9 AH TTX or vehicle respectively; Fig 6 B). In contrast, subsequent TTX application caused a significant increase in the % block in large sized neurons in the vehicle group (mean block = 92.2%, SEM 2.7; p <0.0001, Sidak’s post-hoc comparison to vehicle), but not the 4,9 AH TTX group (mean block = 98.8%, SEM 1.3; p = 0.06, Sidak’s post-hoc comparison to 4,9 AH TTX).

**Figure 6.**
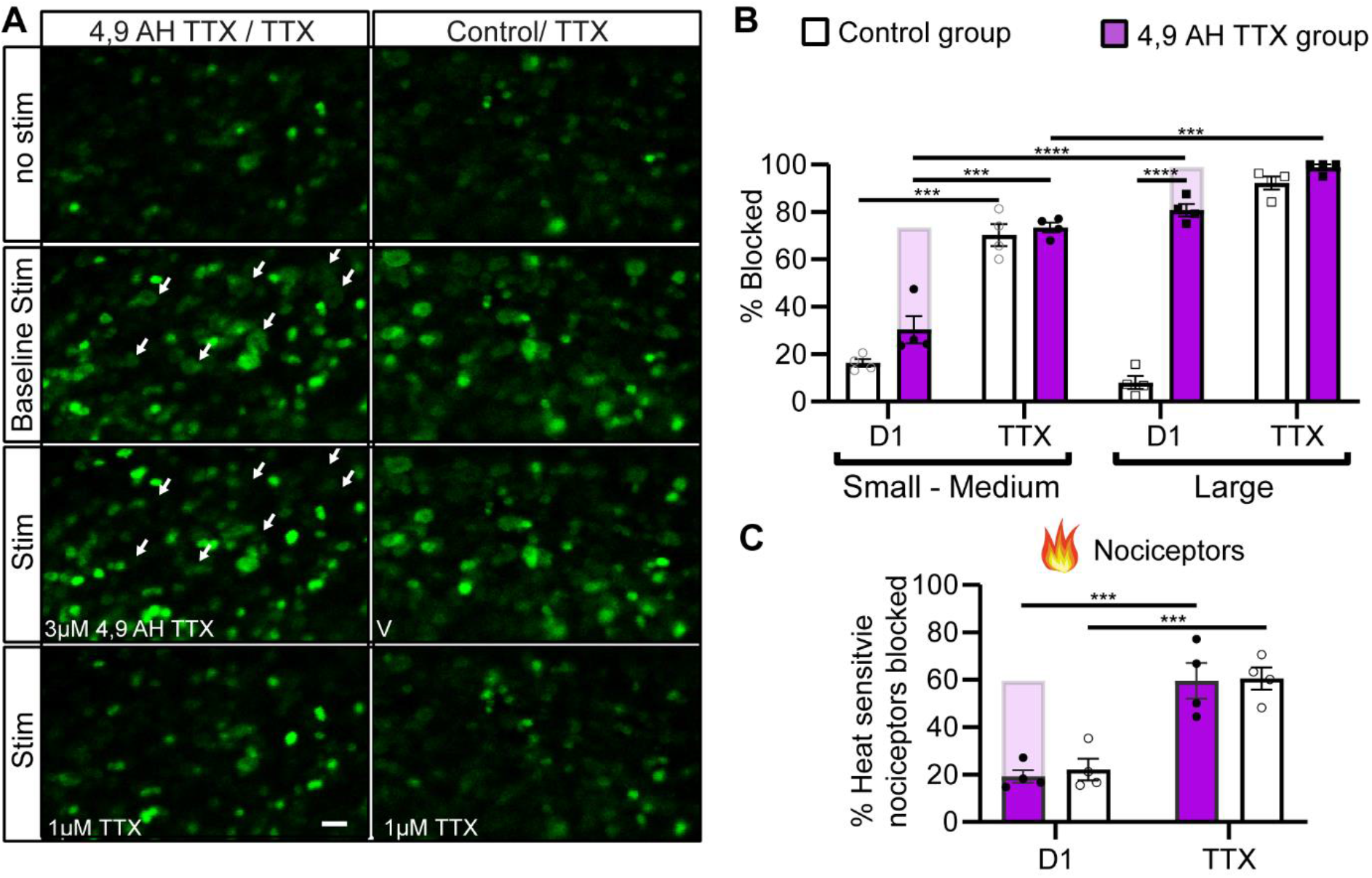
Axonal 4,9 AH TTX application blocked propagation in most large sized neurons but few small to medium sized neurons and very few nociceptors. Example images of L4 DRG cell bodies from in-vivo GCaMP6s time lapse recordings (A). Frames were extracted at baseline and at the peak of electrical stimulation at baseline and following each drug dose. Note that nearly all large sized neurons (indicated by white arrows in the left panel) were blocked by 3 μM 4,9 AH TTX, whilst many small to medium fibres are still conducting. The bar graphs show the proportion of neurons blocked following drug application between stimulating and recording site when neurons are grouped by size (B) and for neurons responding to noxious heat stimulation of the terminal (C). Each point represents the proportion of neurons blocked in an individual animal. Between 46-144 small to medium, 15-40 large neurons and 17-52 heat responsive neurons were sampled between animals. The shaded bars represent the proportion of neurons blocked following addition of 1 μM TTX. D1-drug 1 (either 4,9 AH TTX or Vehicle). Bar graphs represent mean ± SEM. Scale bar = 50μM. 10x magnification. 3-way ANOVA for B) and RM 2-way ANOVA for C). ** p<0.01, *** p<0.001, **** p<0.0001 vs control Sidak’s post hoc test. See also Movie 3.

To examine the effects of blocking axonal Nav1.1 and Nav1.6 with 4,9 AH TTX in nociceptors, we compared block in neurons that responded to noxious heat at the beginning of the experiment. In contrast to the % block in small to medium sized neurons, a lower proportion of heat sensitive nociceptors were blocked by 4,9 AH TTX (19.2%, SEM, 2.7), which was not significantly different to the control (22.1%, SEM, 4.6; F (1, 6) [Drug] = 0.08, P=0.77, 2-WAY RM ANOVA). Subsequent addition of TTX to the sciatic nerve had a significant effect on axonal propagation in heat sensitive nociceptors (F (1, 6) [TTX addition] = 137.8, p<0.0001). The proportion of heat nociceptors blocked by TTX increased to 59.6% (SEM 7.5) in the 4,9 AH TTX group and 60.5% (SEM 4.7) in the vehicle group, which were both significantly different to the block produced previously (both p<0.001, Sidak’s post-hoc comparison to block following 4,9 AH TTX or vehicle respectively; Fig 6 C).

## Discussion

The proportion of C-fibre nociceptive neurons that depend on TTX-S Navs for axonal propagation between the peripheral terminals and the DRG is reported to vary between studies [27; 37; 52], and the contribution of Nav1.7 in particular has not been well defined. In this study, we utilised blockers of Nav1.7 and other TTX-S channels to determine the precise contribution of Nav1.7 to axonal propagation in nociceptors. Specifically, we have shown that only around 2/3 of nociceptors innervating the skin, and less than half of those innervating the muscle, depend on TTX-S channels for axonal propagation. A similar proportion of C-fibre nociceptors innervating the skin were blocked using a selective inhibitor of Nav1.7, suggesting that this channel is the most essential for propagation in the TTX-S population. Finally, our data has revealed that several polymodal mechanoheat sensitive and TRPV1 expressing nociceptors innervating the skin are not dependent on TTX-S Navs for propagation.

Around 2/3^rds^ of nociceptive sensory neurons innervating the skin were blocked following axonal application of TTX. While some previous studies examining TTX block of C-fibre CAPs have reported up to 96% block of propagation along peripheral nerve [27; 37], it is possible that TTX block in these studies was overestimated due to temporal dispersion of compound C-potentials. Moreover, these studies examined C-fibres CAPS in peripheral nerves that contain sympathetic C-fibres, which express Nav1.7 but not TTX-R channels [55], potentially confounding interpretation. Our electrophysiological data revealed that, similar to TTX block, 2/3^rds^ of C-fibre neurons were blocked by axonal application of the Nav1.7 inhibitor PF007, which suggests that Nav1.7 is the TTX-S Nav subtype essential for axonal propagation in this population of nociceptors. However, this does not necessarily mean that Nav1.8, which is expressed in virtually all nociceptors [44; 55] and the major contributor to the upstroke of the nociceptor action potential [38], has no role in propagation in this population. For example, it is plausible that a block of Nav1.7 prevents the membrane reaching the more depolarised threshold required for Nav1.8 activation.

It is not clear why PF007 failed to reproduce the same level of propagation block when examined in our in vivo calcium imaging experiments. Mouse strain differences can be ruled out because a similar proportion of C-fibre neurons were blocked by PF007 ex *vivo* in c57 mice compared to that found in vivo in CD1 mice. Although different forms of electrical stimulation were used to assess axonal propagation between experiments (imaging 4Hz vs 0.1-0.2Hz electrophysiology), higher frequency stimulation would have been expected to facilitate block based on the state dependent characteristics of PF007 [1].

Our electrophysiological and calcium imaging data revealed that ~1/3rd of nociceptors do not depend on Nav1.7 or other TTX-S Navs for axonal propagation. This might explain why peripherally restricted Nav1.7 inhibitors have not provided effective pain relief for chronic pain suffers [25]. It also suggests that there is a sizable population of neurons in which Nav1.7 and other TTX-S Navs are not required for membrane depolarization in order to generate action potentials. As TTX-R neurons were blocked by 1mM lidocaine, which is likely only selective for Navs [42; 46; 53], axonal propagation in these neurons is dependent on the only other TTX-R Nav’s expressed in sensory neurons, Nav1.8 and Nav1.9. Indeed, a previous study identified Nav1.8 as the essential TTX-R channel for propagation in this population [27]. We did not attempt to repeat this finding via pharmacological Nav1.8 inhibition because the selectivity of most Nav1.8 compounds against Nav1.9 have not been tested due to the difficulty of expressing Nav1.9 channels in heterologous cell lines [32], and other selective compounds, such as A-803467, are reported to lack efficacy in rodent peripheral nerve preparations [27]. Nonetheless, our findings provide further indirect evidence in support of Nav1.8 being the most critical Nav in the TTX-R population: we found that many TTX-R neurons were TRPV1 peptidergic nociceptors, which are reported to lack Nav1.9 protein [12; 40].

It is reported that almost every nociceptor (~90%) expresses Nav1.8 protein [44]; this raises the question of why some nociceptors, but not others, can propagate action potentials without other Navs. Although the relatively depolarised threshold for Nav1.8 activation implies that other Navs are usually required for action potential generation [38], reports demonstrate that some nociceptors can generate [47] and propagate [27] signals with Nav1.8 channels alone, and this suggests that Nav1.8 channels in these neurons could have unique gating properties to enable this. For instance, they may have different post-translational modifications, or contain different beta subunits combinations [22], which causes channels to activate at more hyperpolarised potentials, thereby enabling function without other Navs. Alternatively, this population of neurons could have a more depolarised resting membrane potential [11], which would negate the need for other threshold Navs.

Our in vivo imaging data revealed that polymodality is common in sensory neurons innervating glabrous skin, which is consistent with that previously found in our lab using this technique [8]. We found that a higher proportion of small to medium sized M-H polymodal (~40%) were resistant to axonal TTX compared to those only sensitive to mechanical stimuli, which likely reflects that some of the neurons that are exclusively mechanosensitive are low threshold non-nociceptive neurons that don’t typically express TTX-R Navs. The proportion of polymodal neurons blocked by TTX along the axon are similar to that reported previously when examining TTX-R transduction at the peripheral terminals [47; 57]. Although it is tempting to speculate that the neurons resistant to TTX at the terminal represent an overlapping population to those TTX-R along the axon, TTX is reported to differentially block excitability at the terminals compared to the axon in jugular nociceptors, which suggests the expression of TTX-R Navs may differ between neuronal compartments in some neurons [28].

Most high threshold pinch responsive muscle afferents (55%) were not blocked by axonal TTX application, whereas all low threshold cutaneous afferents were blocked by TTX, suggesting that full block of TTX-S channels was achieved and that these propagating muscle afferents do not depend on TTX-S Navs for propagation. Our data are consistent with previous reports that numerous high threshold C-fibre nociceptors innervating the muscle do not require TTX-S Navs for action potential propagation [31; 45]. The TTX-R muscle afferents were blocked by subsequent application of lidocaine and therefore depend on TTX-R Navs for propagation. The hallmarks of chronic pain - ectopic activity in nociceptors, spontaneous pain and central sensitisation within the spinal cord - are more likely to arise following injury to the nerves and tissue of deep structures, compared to those of the skin [26; 49; 54]. Therefore, preferential targeting of Nav subtypes expressed in muscle afferents could be an effective strategy for blocking abnormal activity in these neurons. Given that many muscle nociceptors do not depend on Nav1.7 for propagation, and that pain that is deep in origin is a frequently reported in many chronic pain conditions [2; 4; 30; 34], it is unlikely that peripherally restricted Nav1.7 inhibitors would provide effective pain relief in these individuals.

The TTX block of nearly all large sized neurons or those that responded to low threshold brush suggests that large sized low threshold responsive afferents depend on TTX-S channels for axonal propagation, which is consistent with previous findings [27; 37; 43; 52]. Our electrophysiological data revealed that rapidly conducting A-fibre neurons (likely consistent with the large sized neurons imaged) were not blocked by Nav1.7 inhibition, thereby suggesting that the channel does not make a substantial contribution to propagation in these neurons. In contrast, nearly all the large sized neurons were blocked with 4,9 AH TTX, a selective inhibitor of Nav1.1 and Nav1.6, suggesting that either one or both channels support axonal propagation in this population. Nav1.6 is expressed at nodes of Ranvier in large sized neurons [5; 6], and a previous pharmacological study identified it as being the major contributor to axonal propagation in A-fibres [52].

## Conclusion

Our data indicate that Nav1.7 has an essential role in axonal propagation between cutaneous terminals and the DRG in only around 2/3rds of nociceptive neurons, whereas propagation in the remaining third appears critically dependent on TTX-R Nav subtypes. In the majority of muscle afferent nociceptors propagation is not critically dependent on Nav1.7 or other TTX-S channels.

## Acknowledgments

This research was supported by a research grant from Mundipharma Research Limited, Cambridge, UK. We thank Professor Thomas Sears and Dr Martyn Jones for helping to set up in vivo electrophysiological experiments.

## Author contributions

G.G. conceived, designed, and performed all the experiments, analysed data and wrote the manuscript; S.M conceived experiments and provided conceptual input on the project. F.D. provided conceptual input on the manuscript and corrected the manuscript; E.B.S provided conceptual input on the project and corrected the manuscript; S.B.M. conceived, designed, supervised, and corrected the manuscript.

